# Single-cell alternative polyadenylation analysis reveals mechanistic insights of COVID-19-associated neurological and psychiatric effects

**DOI:** 10.1101/2025.05.02.651855

**Authors:** Qun Chen, Ying Gu, Shuai Liu, Xingyu Li, Ruizhi Xu, Ruixi Ye, Jingjing Yang, Wanshan Ning

**Affiliations:** Institute for Clinical Medical Research, The First Affiliated Hospital of Xiamen University, School of Medicine, Xiamen University, Xiamen, Fujian 361003, China; Xiamen Cell Therapy Research Center, The First Affiliated Hospital of Xiamen University, School of Medicine, Xiamen University, Xiamen, Fujian 361003, China; Department of Pulmonary and Critical Care Medicine, the First Affiliated Hospital of Xiamen University, School of Medicine, Xiamen University, Xiamen, Fujian 361003, China

**Keywords:** COVID-19-associated neurological and psychiatric effects, single-cell APA analysis, neural cell, functional APA event, disease risk gene

## Abstract

COVID-19 is associated with increased risks of neurological and psychiatric sequelae. Alternative polyadenylation (APA) is ubiquitous in human genes, resulting in mRNA diversity, and has been validated to play a pivotal regulatory role in the onset and progression of a variety of diseases, including viral infections. Here, we analyzed the APA usage across different cell types in frontal cortex cells from non-viral control group and COVID-19 patients, and identified functionally related APA events in COVID-19. According to our study, the poly(A) site (PAS) usage is different among cell types and following SARS-COV-2 infection. Moreover, we found the genes with significant PAS level changes affected pathways related to RNA splicing, and neuronal development and function, suggesting that survivors of COVID-19 will have a high risk of these diseases and that alternative splicing functions cause these changes. Additionally, APA usage and its correlation with gene expression levels varied across genes, some prefer short isoform that is more stable to produce more proteins, while others may be regulated by different mechanisms. A total of 267 risk genes targeted by microRNAs for common neurological and psychiatric disorders were found to undergo significant changes in APA following infection. In conclusion, our comprehensive analysis of APA in neural cells from COVID-19 patients at the single-cell level elucidated changes in APA levels in the brains of SARS-COV-2-infected patients and confirmed that these changes impair the function of the nervous system, providing important insights for COVID-19-associated sequelae.

**Highlights:**

1. We analyzed the changes in single-cell APA level in the frontal cortex after SARS-COV-2 infection and found that PAS usage is different among cell types and changed after infection.
2. We find that APA changes are related to RNA splicing, and neuronal development and function, such as synaptic plasticity, nervous system development, learning or memory and cognition, and the brain function may be impaired due to global gene expression and APA changes.
3. A total of 267 risk genes for common neurological and psychiatric disorders were found to undergo significant changes in APA following infection, implying that APA alterations in specific genes could be a potential therapeutic target and predictive biomarker for neurological and psychiatric disease.
4. APA in infection samples results in the removal/addition of miRNA target sites in the 3′-UTR of genes. A total of 2404 miRNAs were predicted which suggests that the miRNAs’ target site information can indicate the impact of APA events and has the potential to be used as a predictive biomarker.

## Introduction

Brain and peripheral nerve tissues express a larger proportion of the genome than any other tissue, resulting in high levels of mRNA diversity^[1]^. This diversity is largely due to alternative splicing, transcription initiation sites, and polyadenylation^[2]^. APA, which occurs in more than 70% of human protein coding genes, has recently been considered to be a key regulator of gene expression during disease onset and progression^[3]^. Moreover, the studies have shown that compared to other organs/tissues, the genes expressed in the nervous system contain relatively longer 3′-UTRs on average^[4]^. Among them, the hippocampus shows the largest number and expression of 3′-UTR extension in the nervous system^[1]^. The hippocampus plays a role in memory and spatial localization. Cerebral hypoxia and encephalitis can lead to hippocampal injury and memory loss. The hippocampus of mice suffering from status epilepticus and epilepsy, more than 25% of the transcriptome shows changes in the length of poly (A) tail ^[5]^. The de-adenylation disproportionately affects genes previously associated with epilepsy^[5]^.

COVID-19 survivors are at the risk of long-term neurological diseases. The study found that 14% of the patients are detected with new cerebral ischemic lesions and 86% of patients have astrocyte proliferation, microglia activation and cytotoxic T lymphocyte infiltration in the brainstem and cerebellum^[6]^. In addition, COVID-19 is associated with a severe innate immune response and systemic inflammation to promote cognitive decline and neurodegenerative diseases, which makes COVID-19 survivors likely to suffer from neurodegenerative diseases in the next few years ^[7]^. Studies have shown that polyadenylation plays an important role in the behavior of the immune system. For instance, the inhibition of polyadenylation reduces the induction of inflammatory genes, and translation is inhibited by regulating the cellular polyadenylation binding protein in inflammatory response, emphasizing the necessity of studying the regulation of polyadenylation in the immune system^[8-10]^. The main resident immune cells of the brain are microglia, a large number of perivascular macrophages and dendritic cells, and detectable numbers of T cells, B cells, and natural killer (NK) cells. In addition, the average 3′-UTR length shortens after vesicular stomatitis virus infection in macrophages^[11]^.

To the best of our knowledge, the APA regulation in association with COVID-19 has not been studied in brain yet. In view of the extensive regulatory role of APA in neurons and immune response, the study of APA level in COVID-19 patients and survivors’ brain tissue is helpful to understand the pathogenic mechanism of SARS-CoV-2 from the level of post transcriptional modification, which is of great significance to the nerve injury in COVID-19 patients.

## Materials and Methods

### Data Pre-processing and snRNA-seq Quality Control

The data are public available and deposited in the GEO with accession number GSE159812 by 3’ tag-based sequencing protocol 10x Genomics v3^[12]^. The 15 samples of brain frontal cortex from non-viral group and COVID-19 group are chosen. Raw gene counts are aligning reads to the hg38 genome (refdata-gex-GRCh38-2020-A) by CellRanger software (V6.0) (10x Genomics). The cells with high ratio of mitochondrial (more than 10%) and the features less than 200 are treated as the outliers and removed in Seurat 4.0.5^[13]^.

### Cells Integration and Annotations

After quality control, the data are further normalized, scaled, and filtered the highly variable data features for principal component analysis (PCA) by Seurat functions of NormalizeData, ScaleData and FindVariableFeartures. Then 15 samples are integrated by Harmony method to reduce the individual difference. The entire dataset is projected onto two-dimensional space by UMAP (Uniform manifold approximation and projection) on the top 20 principal components following the tutorial of Seurat. We annotate cell types referring to the original paper^[12]^ at gene expression levels.

### Gene Set Enrichment Analysis

We use irGSEA(v.4.0.0) to analyze and comprehensively evaluate the differential gene sets for cell subsets in the control group and the COVID-19 group. Firstly, we scored individual cells by multiple gene set enrichment methods, and generated multiple gene set enrichment score matrix. Next, we calculated the differentially expressed gene sets for each cell subpopulation in the enrichment score matrix of each gene set by wilcox test. Then the robust rank aggregation (RRA) algorithm in RobustRankAggreg package is used to comprehensively evaluate the results of difference analysis, and the gene sets that are significantly enriched in most of the gene set enrichment analysis methods are screened, visualized and analyzed.

### Cells Annotations at APA Level

According to the tutorial from https://github.com/YangLab/SCAPTURE^[14]^, after the raw data processing following the method described in chapter 3, we get the matrix with PAS transcripts files as the input for single-cell APA analysis, (the process is consistent with single cell gene expression for filtering outliers, normalization, integration, and clustering). The clusters are annotated with the expression of the gene transcripts with specific PAS.

### APA Usage Calculation

The preference of proximal PAS usage between single cells, the mean value of proximal PAS usage across expressed polyadenylated genes per cell is determined by the SCAPTURE. From the SCAPTURE manual, we choose the PUI (proximal usage index) method as the index of APA usage preference. It calculates the ratio of counts per million (CPM) with proximal usage to the CPM with total PAS usage. The usage levels are ranging from zero to one. For the PAS transcripts with high PUI, it indicates that this PAS transcript prefers to be used at the proximal location, while the PAS transcripts with low PUI, the cells are prone to choose the distal PAS. Each transcript with identified PAS has this value to show their APA usage preference.

### The GO Analysis and KEGG Analysis Based on the APA

All the subtypes of cell clusters are extracted from the APA usage matrix, and the differential expressions are calculated. Transcripts with p values less than 0.05 and an average log2 fold change larger than 0.25 are selected after the data is obtained for differential analysis. The multiple transcripts with distinct PAS transcripts could be produced from a single-gene locus, and we use the genes with significant changes of PAS transcripts as the input for Gene Oncology (GO) biological function and Kyoto Encyclopedia of Genes and Genomes (KEGG) pathway. GO process and pathway enrichment analysis of genes with significant alterations of PAS were performed by metascape^[15]^. The similarity score between terms was determined by calculating the Kappa-test scores between each of the two representative terms chosen from the Metascape results (P value ≤ 0.05). Cytoscape^[16]^ was employed to visualize the representative GO terms, and terms were set as nodes, and similarity scores of greater than 0.3 were set as edges.

### The Validation of PAS Usage by IGV

Reads from each cell type are extracted from the bam file, and the bam file is converted to the binary wig (bw) format file using deeptools. The metadata bed file from SCAPTURE is used as the reference and imported to the IGV software, and the interested cell types of bw file are imported to the IGV for visualization.

### The 3′-UTR Changes in APA

The PAS located in 3′-UTR are extracted and we integrate these transcripts into the movAPA^[17]^ software to calculate the 3′-UTR changes according to its tutorial. For each gene, the ratio of PAS transcript counts in two different groups are calculated at proximal and distal sites by movAPA.

## Results

### Global APA Levels in Neural Cells

The poly(A) sites are annotated using the latest genome annotation, roughly one-third of them were located in 3′ UTRs, while the remaining ones were found in other genomic regions (i.e. 5′ UTR, coding sequence (CDS), and intron) (Figure 1A). The poly(A) signals, which are typically located 15–30 nt upstream of the poly(A) sites, are necessary for pre-mRNA cleavage and polyadenylation^[18]^. The motif enrichment unraveled that the canonical poly(A) signal (AAUAAA) is significantly enriched in the dataset (Figure 1B). Similar to previous studies^[3, 19]^, AAUAAA, AUUAAA, and single nucleotide variants of AAUAAA have been identified as typical poly(A) signals upstream of the poly(A) site (Figure 1C). These results demonstrated the authenticity of poly(A) sites identified by SCAPTURE (Supplementary table 1).

**Figure 1.**
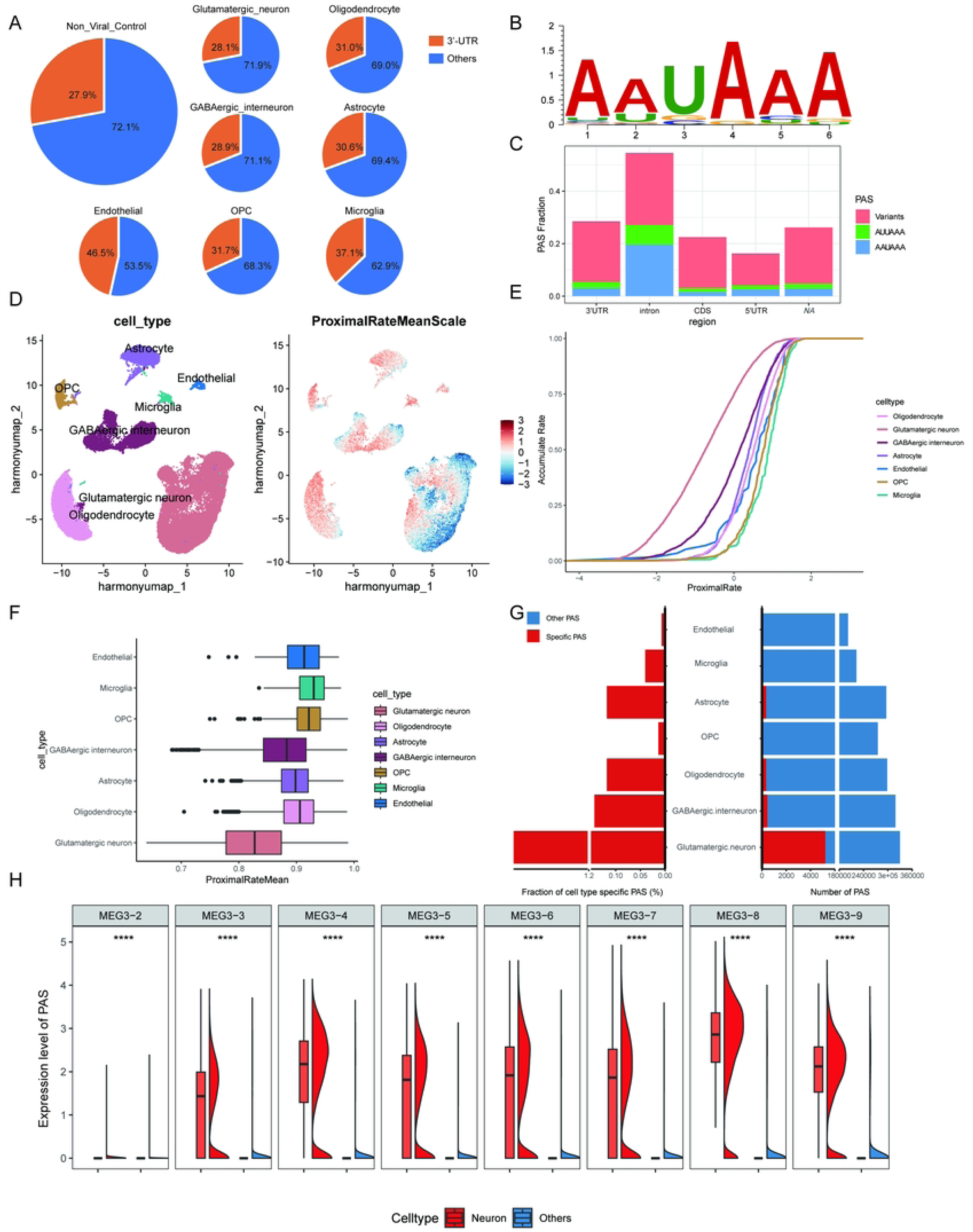
Global APA levels in neural cells. (A) Distribution of poly(A) sites in 3′ UTR. (B) Canonical poly(A) motif (AAUAAA) enrichments for poly(A) sites. (C) Poly(A) signals within the upstream 50 nt of poly(A) sites. (D) Left: UMAP of 16,020 nuclei from the medial frontal cortex of 7 control individuals, identified 7 different clusters. Each point depicting a single cell nucleus, colored according to cluster designation. Right: The mean PAS usage of each single cell for 7 samples after scaling, the expression level is from -3 to 3. (E) Accumulation rate of each cell type based on non-viral control group in solid lines. (F) Proximal rate of each cell type. (G) The numbers of cell-type-specific PAS and their fractions among the total identified PAS in different cell-type populations. (H) Expression levels of MEG3 PAS in neuronal cell types relative to other cell types.

We integrated seven samples of frontal cortex from individuals in the control group, 16,020 nuclei are generated with 24,813 features. Unsupervised clustering identified seven major cell types (Figure 1D), the cell markers that are uniquely expressed in subpopulation refer to the published paper^[12]^. Specifically, glutamatergic neurons are marked by SLC17A7, GABAergic interneurons are marked by GAD1, astrocytes are marked by GFAP, oligodendrocytes are marked by ST18, oligodendrocyte progenitor cell (OPC) is marked by VCAN, microglia are marked by CD74 and endothelial cells by CLDN5. Supplementary Figure 1A shows that the glutamatergic neurons are the largest population, oligodendrocytes, and GABAergic interneurons are higher than other cell types, and the number of endothelial cells and microglial is small. The top 10 most enriched genes in each cluster are shown in the heatmap of Supplementary Figure 1B and the violin plot shows the differential expression of cell markers with significantly different percentages per cell type (Supplementary Figure 1C, Supplementary table 2).

For glutamatergic neurons, the up-regulated gene sets are enriched in Alzheimer’s disease (AD), and amyotrophic lateral sclerosis, belonging to calcium ion signaling and other related pathways, and conversely, down-regulated genes in astrocytes, OPC, oligodendrocytes, microglia, and endothelial cells are also enriched in these pathways. For GABAergic interneurons, genes are mainly enriched in metabolic pathways, the up-regulated genes are enriched in β-alanine, aspartate-alanine-glutamic acid and butanoate metabolism, and citrate cycle, and down-regulated in unsaturated fatty acid biosynthesis and adherens junction. For microglia, genes enriched in antigen processing and presenting are activated. These cells are more likely to participate in inflammation-related responses after infection (Supplementary Figure 1D).

We calculated the APA usage in frontal cortex cells, the average PUI of each cell is projected into each cell type based on the gene expression in UMAP shown in Figure 1D. The pink color represents the scaled proximal rate mean of each cell, and the glutamatergic neurons and GABAergic interneurons have a relatively deep blue color referring to the mapping of cell subtypes, which indicates they prefer to choose the non-proximal PAS. These results are consistent with the previous reports that the neurons prefer to choose the distal PAS^[2, 20]^. Subsequently, we described the accumulation plot and box plot of each cell type to further systematize the integrated APA usage, and we noticed that the average PUI of neurons is less than that of other cell types (Figure 1E, F). Furthermore, of all the PAS identified in various cell-type populations, neurons account for the greatest number as well as the largest fraction of cell-type-specific PAS (Figure 1G). So we pay more attention on changes in PAS levels in neurons, compared to other neural cells.

To address the functional consequences of APA events, we tested for the enrichment of gene ontology and functional pathway annotations among the genes with differential PAS used in neurons (excitatory neurons and inhibitory neurons) compared to non-neuronal nerve cells (Supplementary table 3). Not surprisingly, these genes were highly enriched in biological processes relating to neural development or neural function (Supplementary Figure 2). In summary, we observed that a set of genes with neuronal regulatory functions was preferentially subject to distal PAS in neurons.

Neuronal cell types exhibited significantly higher expression of MEG3 PAS. MEG3 has been related to various processes such as necroptosis, apoptosis, inflammation, oxidative stress, endoplasmic reticulum stress, and epithelial-mesenchymal transition, according to recent studies^[21-23]^. In neurons, the usage of PAS sites was significantly higher in different MEG3 transcripts than in other neural cells (Figure 1H).

### Changes in Global APA Levels in Neural Cells Following Infection With SARS-COV-2

The control group (Figure 2A) and positive group (Figure 2C) display different patterns of preference, the shift to proximal in neurons is obvious after being infected with SARS-CoV-2, and it indicates the wide changes in APA levels after infections. Each cell type proportion in the non-viral control group, COVID-19 group, and total are calculated. The excitatory neurons are the largest population; oligodendrocytes and inhibitory neurons are larger than other cell types, and the number of endothelial cells and microglials are small (Figure 2B). The accumulation plot of the infected group described that the neurons’ average PUI is less than other cell types, but the difference becomes smaller after infections (Figure 1E, Figure 2D).

**Figure 2.**
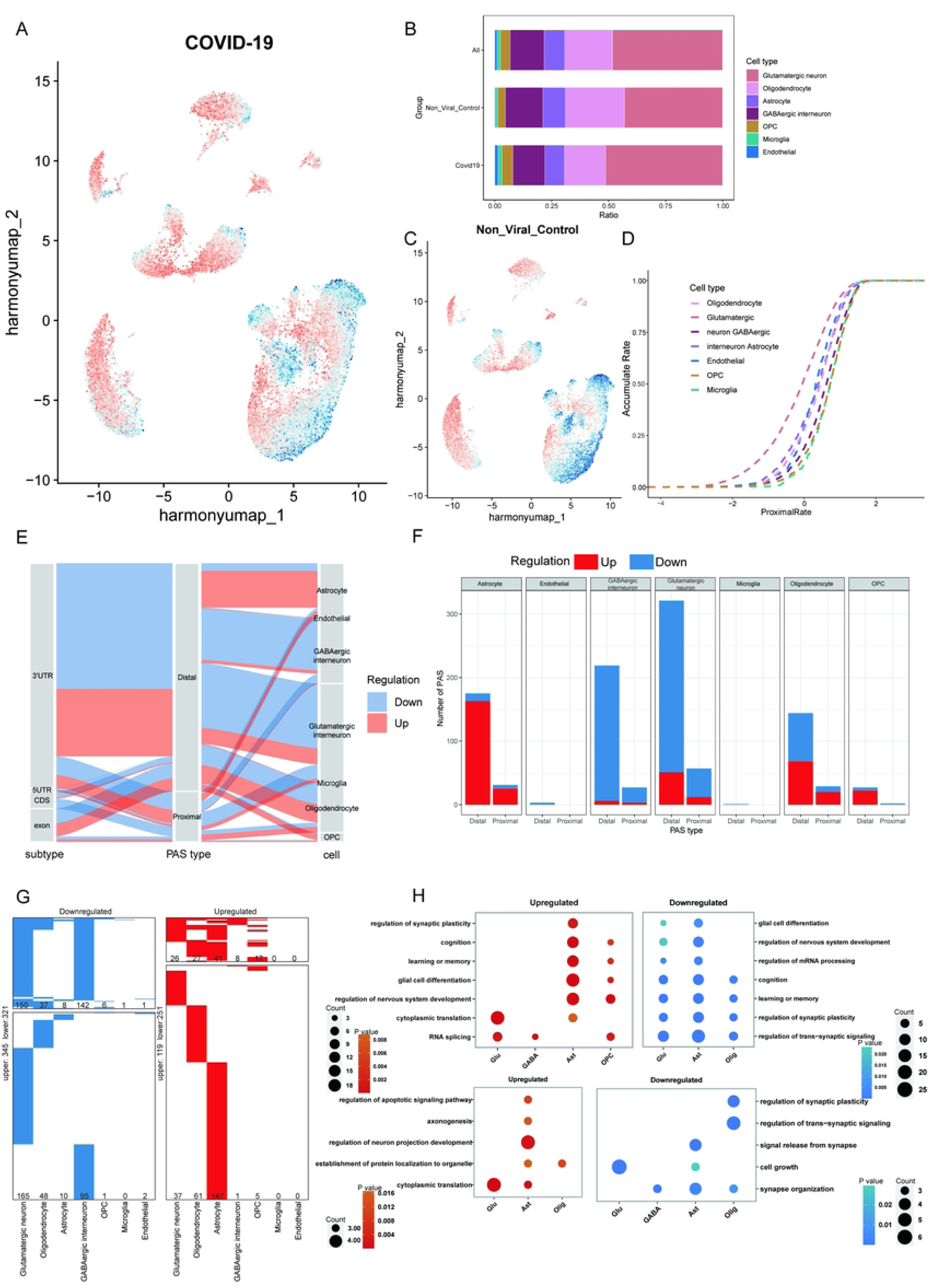
Changes in global APA levels in neuronal cells following infection with SARS-COV-2. (A) The global preference of APA usage in the infected group. (B) Each cell type proportion in the non-viral control group, COVID-19 group, and total. (C) The global preference of APA usage in the non-viral control group. (D) Accumulation rate of each cell type based on the infected group with dashed lines. (E) Changes among the PAS location, PAS type, and cell types, the pink color represents the upregulated PAS changes, and the blue represents the downregulated PAS changes. (F) The count number changes of PAS in different cell types after infection, the pink color represents the upregulated PAS, and the blue color represents the downregulated PAS. (G) Heatmaps showing the distribution of upregulated (red) and downregulated (blue) PAS for each cell type in the brain between the non-viral control group and the infected group. PAS not differentially expressed are in white and the numbers of PAS are indicated. The upper part indicates the PAS shared by at least two cell types, and the lower panel indicates the unique PAS of each cell type. The numbers of PAS are annotated on the plots. (H) Dot plot showing the representative GO terms enriched for distal and proximal PAS in different cell types. The upper part indicates the distal PAS, and the lower panel indicates the proximal ones.

Based on the above results, the details of APA changes in each cell cluster are explored and the significance of changes is assessed by Kolmogorov-Smirnov test. For the different nerve cell types, mainly glutamatergic neurons and GABAergic neurons, the PAS preferences are consistent and have a relatively high significant preference for the proximal usage of PAS (Supplementary Figure 3A, 3B). We find a slight tendency to shift to the non-proximal PAS after infection for astrocytes (Supplementary Figure 3C), oligodendrocytes (Supplementary Figure 3D), OPC (Supplementary Figure 3E), microglia (Supplementary Figure 3F), and Endothelial (Supplementary Figure 3G).

The different trends after infection may indicate the APA usage changes among these cell types. Subsequently, we investigated the detailed dynamic changes of each PAS after infections, we compared the infected group with the control group and used the Sankey plot to show the flow dynamic. The left grey column represents the PAS location in the exon of genes (Figure 2E), and the 3′-UTR accounts for the largest proportion. The second column is the PAS type, and we treat the non-proximal PAS as the distal proximal PAS here to better calculate the PAS type. The distal proportion is larger than the proximal PAS, which indicates the cells in brain tissues prefer to use the distal PAS instead of the proximal PAS. The third column is the cell type. The ratio of proximal PAS to distal PAS and specific counts in different cell types were calculated by bar stacking plots (Figure 2F). It is obvious that neurons, including glutamatergic neurons and GABAergic interneurons, exhibited downregulated PAS (in blue color), whereas certain PAS were upregulated following infections. In astrocytes, oligodendrocytes, and OPC, the upregulation trend (in pink) accounted for a larger proportion of the cells, suggesting a tendency for these cell types to have reduced proximal PAS use compared to the non-viral controls (Supplementary table 4).

The data in Figure 2E and Figure 2F are consistent with the accumulation rate in the proximal rate shift in Supplementary Figure 3. In these two types of neurons, the increased proportion of downregulation PAS in proximal sites (Figure 2F) results in the smaller mean PUI after infections, which could explain why neurons have an increasing proximal accumulation rate after infection in Supplementary Figure 3A, 3B. Besides, the other types of cells have a relatively low proportion of downregulated PAS in proximal sites and more upregulated PAS in distal sites, and the proximal accumulation rates are decreasing in these cell types (Supplementary Figure 3C-G).

To validate the analysis results, we used the raw bam file from Cellranger, split the bam file by cell types, and transferred the bam file to the wig file. Then we use the IGV to browse the proximal and distal usages. Here are the validation results of some PAS in Supplementary Figure 4. We choose the different PAS expressions between the infected and control groups from each cell type, the first four samples from two groups are used here. We use the peak height in sample tracks to indicate the expression level of PAS and compare the results after the SCAPTURE calculation.

The trends of PAS changes in these six cell types are consistent with the results shown in supplementary table 5. Take oligodendrocytes as an example, PAS sites at MT3 gene (Supplementary Figure 4D), we find the height of tracks in the control group is higher than infected group, it is consistent with the result in supplementary table 5. Compared with the control group, the infected group shows down PAS regulation at the MT3 gene, the average log2 fold change is -1.1932.

To further investigate the effects of these significant PAS site usage at the cellular level, we compared the PAS expression of cell types between groups. To this end, we identified more than one thousand differentially expressed PAS (|avg_logFC| > 0.25) in at least one cell type. 370 upregulated and 666 downregulated PAS were identified. Notably, approximately half of the differentially expressed PAS were specifically detectable in the neurons and astrocytes (Figure 2G).

Gene ontology (GO) analysis of differentially expressed PAS further uncovered that downregulated distal PAS in glutamatergic neurons was enriched largely in neuronal development and function (Figure 2H), such as synaptic plasticity, nervous system development, learning or memory, cognition, and so on, highlighting the importance of downregulated expressed distal PAS in neurons after infection, while downregulated expressed proximal PAS in astrocyte and oligodendrocyte were mainly associated with synaptic function after infection.

### The APA Usage and Its Correlation with Gene Expression Level

It has been reported that the choice of PAS could regulate the post-transcription and change the gene expression levels. We utilize the glutamatergic neurons and GABAergic interneurons to investigate the correlation between APA usage at 3′-UTR and gene expression level. To explore this correlation, movAPA was employed to calculate the different PAS usage between the COVID-19 group and the control group. We get the different usage values by the value of the infected group to subtract the usage value of the normal group, and define the absolute value larger than 0.1 as the significant changes of APA usage. Combined with gene expression, we could get four kinds of information in the four quadrants of Figure 3A and Figure 3B. In the first quadrant, APA usage value of the infected group is relatively larger, and the corresponding gene expression is increased; In the second quadrant, the APA usage value of the infected group is relatively small, and gene expression is increased; In the third quadrant, the APA usage value of the infected group was relatively small, and the gene expression was decreased; while in the fourth quadrant, both APA usage and gene expression are decreased.

**Figure 3.**
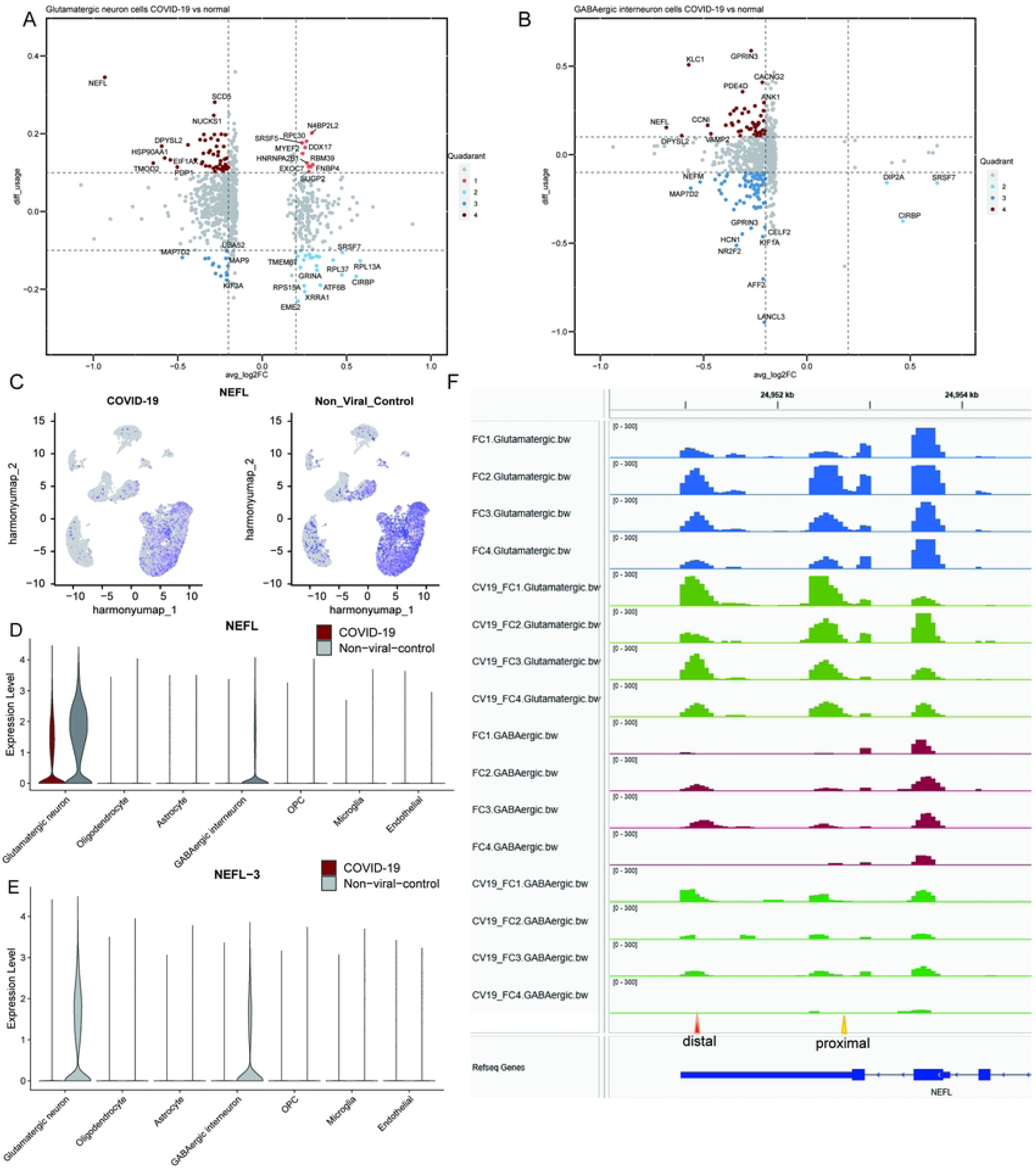
The APA usage and its correlation with gene expression level. (A) APA usage and its correlation with gene expression level of glutamatergic neurons and (B) GABAergic interneurons. (C) Global gene expression of NEFL in the infected (left) and control (right) group. (D) NEFL gene expression in cell clusters. (E) NEFL-3 APA levels in cell clusters. (F) The PAS usage of NEFL at distal and proximal sites by IGV.

In glutamatergic neurons (Figure 3A), the genes DDX17, RBM39, FNBP4, and others are located in the first quadrant indicating these genes prefer to use the distal PAS sites and the gene expression level increased. Genes like RPL37, RPL13A, GRINA, ATFB, and CIRBP located in the second quadrant prefer to use the proximal sites with increased gene expression. As for MAP7D2, MAP9 or others in the third quadrant tend to use the proximal sites, and gene expression is relatively downregulated in the infected group. The genes located in the fourth quadrant like NEFL, SCD5, and HSP90AA1 prefer to use distal PAS sites with the downregulated gene expression level. For GABAergic interneurons in Figure 3B, we find that some genes have similar location patterns. For example, CIRBP is located in the second quadrant in these two types of cells, and NEFL in the fourth quadrant, for both cell types. For other cell types, NEFL is also predominantly located in the fourth quadrant (Supplementary Figure 5, Supplementary table 6).

NEFL is a light chain of neurofilaments that functionally maintain the neuronal caliber. Individuals with dementia have elevated levels of NEFL in the cerebrospinal fluid (CSF) ^[24, 25]^. Based on the gene expression level in Figure 3C, the NEFL is mainly enriched in glutamatergic neurons and GABAergic interneurons, the NEFL expression level is decreased obviously for the COVID-19 group in UMAP. The NEFL expression level of each cell cluster is shown in Figure 3D. For each cell cluster’s APA usage level shown in Figure 3E, the PAS site usage in NEFL-3 is decreased after infection.

To validate the result of NEFL, we visualized the abundance of PAS sites using wig file of excitatory neurons and inhibitory neurons obtained from raw bam file by IGV shown in Figure 3F.

The gene NEFL is located in the chromosome negative strand, and the narrow bar represents untranslated region. Thus, the 5′-UTR and 3′-UTR are located in the right and left side respectively. We use the orange triangle at the bottom to indicate the proximal site, and red triangle to indicate the distal site. Here, the average peak heights from case group in distal sites to the total sites are higher than the control group, especially in sample CV19-FC3 of Figure 3F. This result is consistent with Figure 3A-B, that is, NEFL prefers to use the distal PAS sites after infection.

### Genes with Variational APA Levels and Differential Expression Widely Affected Neural Function in MicroRNA-dependent Fashion

Since the previous study demonstrated the pathogenic role of APA events in a microRNA-dependent fashion, this result suggests that the microRNAs (miRNAs) target site information can indicate the impact of APA events and has the potential to be used as a predictive biomarker^[26, 27]^. APA in infection samples results in the removal/addition of miRNA target sites in the 3′-UTR of genes.

Subsequently, a total of 2404 miRNAs (Supplementary table 7) targeting the 408 genes in the four quadrants of all cell types were predicted using the miRcords^[28]^, miRTarBase^[29]^, and TarBase^[30]^. Nearly half of them were cell-type-specific (Figure 4C, 4D).

**Figure 4.**
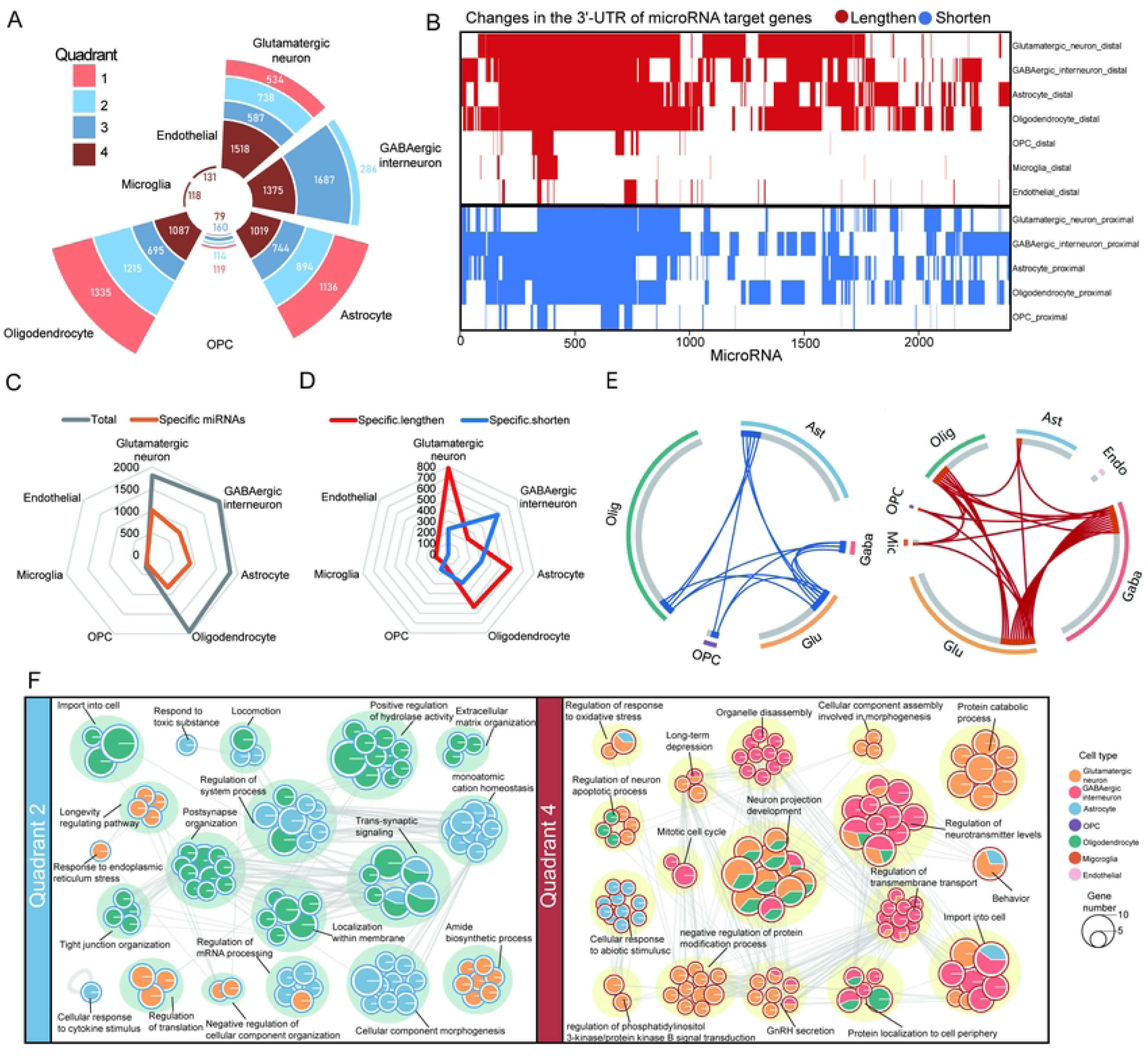
Genes with variational APA levels and differential expression widely affected neural function in a microRNA-dependent fashion. (A) Radial plot showing the number of microRNAs targeting the four quadrants of genes. (B) Heatmap showing the changes in the 3’-UTR of microRNA target genes, either lengthen (red) or shorten (blue). (C) Radar Charts showing the total, (D) specific microRNAs targeting lengthen or shorten 3’-UTR across cell types. (E) Circos plots show the overlap of genes from different cell types in quadrant 2 (left) and quadrant 4 (right). Connecting lines link the same genes that are shared by cell types. (F) GO terms and pathways of quadrant 2 (left) and quadrant 4 (right) genes in different cell types. The network nodes were displayed as pies. Each pie sector is proportional to the number of hits originating from a gene list. The pie charts are colored by cell types, where the size of a slice represents the percentage of genes under the term that originated from the corresponding cell type. Terms with a similarity score > 0.3 are linked by an edge (the thickness of the edge represents the similarity score).

In order to further investigate the effects of the genes in four groups, which represents different APA usage and gene expression level, we analyzed these genes across all cell types. In quadrant 2, the largest numbers of genes were observed in Astrocyte and Oligodendrocyte (31 and 51 genes), while glutamatergic and GABAergic interneurons had 50 and 51 genes in quadrant 4 (Figure 4E).

Furthermore, through functional annotation enrichment analysis, we found that “neuron projection development”, “regulation of neurotransmitter levels” were enriched for group 4 in glutamatergic neurons and GABAergic interneurons and Oligodendrocyte, “long-term depression”, “GnRH secretion” in both glutamatergic neurons and GABAergic interneurons. In group 2, “trans-synaptic signaling” and “regulation of system process” were both enriched in Astrocyte and Oligodendrocyte. “RNA splicing”, “glial cell differentiation” and “Parkinson disease” were enriched in group 1 (Figure 4F, Supplementary Figure 6, Supplementary table8).

### Neurological and Psychiatric Disease Risk Genes Are Significantly Altered in APA Following Infection

To unravel the potential mechanisms underlying the development of neurological and psychiatric effects following SARS-COV-2 infection, fifteen common neurological and psychiatric disorders, including AD, Parkinson’s disease (PD), epilepsy, autism spectrum disorders (ASD), depressive disorder, bipolar disorder (BD), attention deficit hyperactivity disorder (ADHD), multiple sclerosis (MS), ischemic stroke, schizophrenia (SZP), amyotrophic lateral sclerosis (ALS), Huntington disease (HD), migraine disorder, brain aging, and anxiety, were selected for joint analysis using the two databases DisGeNET^[31]^, BioKA^[32]^, and diseases from the GWAS^[33]^ catalogue in the published paper^[12]^. After adding up all of the genes in the four quadrants where there were variations in APA levels and differential expression in the seven cell types (glutamatergic neuron, GABAergic interneuron, astrocyte, endothelial, microglia, oligodendrocyte, and OPC), we had a total of 408 genes, 267 of which were identified as risk genes in DisGeNET and BioKA databases and GWAS catalogue (Supplementary table 9)(Figure 5A, 5B).We found that there is a significant correlation between these 267 risk genes and neurological disorders and traits, specifically AD, PD, and schizophrenia. Especially, we observed that CALM1, part of the calcium signal transduction pathway^[34]^, was down-regulated accompanied by a shorter 3’-UTR after infection with SARS-COV-2 (Supplementary Figure 7A, 7B). A recent study finds mice’s dorsal root ganglion development along with hippocampal neuron activation are impaired when *Calm1* long 3’-UTR mRNA isoform eliminated via CRISPR-Cas9 gene editing^[35]^. The amyloid precursor protein (APP) has gained notoriety due to its hypothesized critical function in the development of AD^[36, 37]^. We discovered that expression of APP increased after infection, with oligodendrocytes displaying a longer 3’-UTR and astrocytes a shorter 3’-UTR (Supplementary Figure 7C, 7D). It has been verified that APP 3’UTR length can affect translation efficiencies. A variation in the length of the APP 3’UTR, whether long or short, can cause aberrant expression and eventually increase the risk of AD^[38]^. Actin constitutes a prominent cytoskeletal protein within eukaryotes organisms, and is recognized for its localization in neuronal synapses.

**Figure 5.**
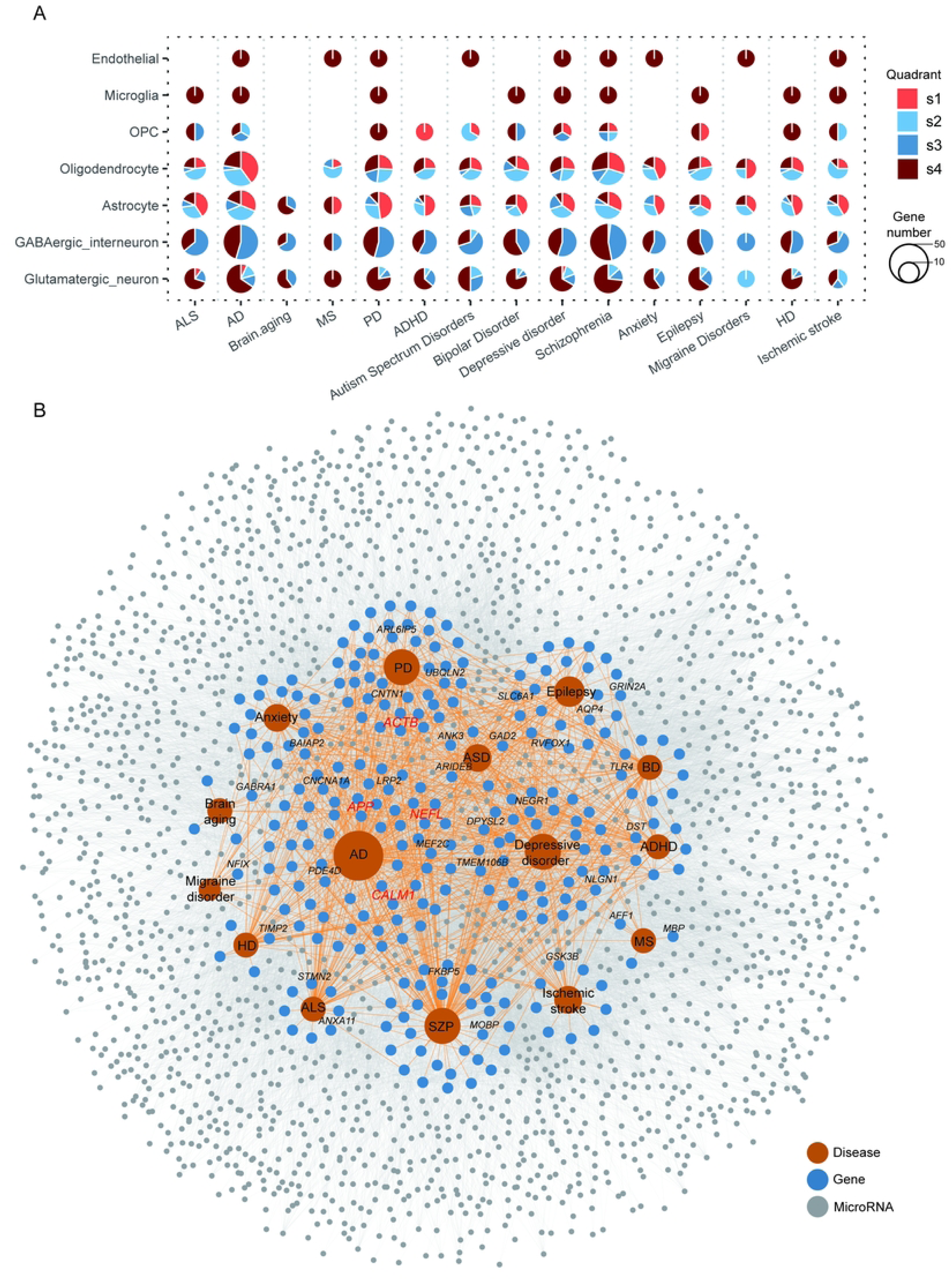
Neurological and psychiatric disease risk genes are significantly altered in APA following infection. (A) Dot plots showing that genes from four quadrants overlapped with genes from neurological and psychiatric disorders associated gene sets. The pie charts are colored by quadrants, with size indicating the frequency of genes. (B)Network visualizing the overlap between the genes in the four quadrants and genes involved in 15 common neurological and psychiatric disorders. The colors of grey, blue, and orange nodes represent targeting miRNAs, genes and diseases, respectively. The colors of grey and light orange lines represent the connection between genes-miRNAs and genes-diseases.

Investagation into beta-actin (ACTB) reveals that two alternative transcripts were induced and terminated at tandem polyA sites, the longer 3’UTR brings higher stability to the transcript, resulting in higher gene expression, in mouse neuronal cells^[39]^. In our study, the expression of ACTB increased after infection, with a longer 3’-UTR in astrocytes (Supplementary Figure 7E, 7F). TMOD2 acts as a fibrogenic gene in idiopathic pulmonary fibrosis (IPF) and exhibits a shortened 3’UTR and increased protein expression levels^[40]^. In the present study, after SARS-COV-2 infection, TMOD2 shows 3’-UTR shortening as well as expression downregulated.

## Discussions

There is growing evidence that SARS-CoV-2 infection causes neurological deficits in a large proportion of infected patients^[6, 41, 42]^. Several studies have reported psychiatric symptoms in patients with COVID-19, and a growing body of research suggests that brain disorders may persist after recovery from primary infection^[43, 44]^.

Though it has been reported that the gene expression level changes after infections^[45, 46]^, the changes in APA level have not been reported in the brain tissue after infection. We select the appropriate method and find significant changes in the global APA levels shown in Figure 2.

With shortened 3′-UTR average length and extensive APA response to viral infection, expression levels of genes with APA are altered and enriched in immune-related pathways, such as Toll-like receptor, RIG-I-like receptor, JAK-STAT, and apoptosis-related signaling pathways^[11]^. In a genome-wide analysis with PBMC RNA-seq dataset of COVID-19 patients^[47]^, APA-related genes are abundant in the ontology classification related to innate immunity, such as neutrophil activation, MAPK cascade regulation and cytokine production, and interferon-γ and innate immune response, suggesting that APA events may be a better predictor of neurological deficits than alternative splicing for COVID-19 patients. Therefore, APA could be used as a potential therapeutic target and a novel biomarker for post-COVID survivors.

In our study, we find that the average utilization of proximal PAS in glutamatergic neurons and GABAergic interneurons increases after COVID-19 infection, indicating that brain cells have a wide range of APA response after SARS-CoV-2 infection, which is similar to the result of APA response caused by other virus infection^[47, 48]^. The selection of PAS is different for various cell subtypes. For neurons, the normal group tends to use distal PAS, which is consistent with previous reports ^[20]^. Both microglia and endothelial cells express a large amount of short 3′-UTR^[20]^. For COVID-19, the excitatory neurons and inhibitory neurons show a preference for distal PAS.

The genes with different APA levels deserve to be studied, especially in the glutamatergic neurons and GABAergic neurons. We extract all PAS in different cell types for further GO analysis to study related gene expression changes following infections, and the impacts of APA levels on altering the overall cellular pathway. We find that APA changes are related to RNA splicing, and neuronal development and function (Figure 2H), such as synaptic plasticity, nervous system development, learning or memory, cognition, and so on. GO analysis shows that brain function may be impaired due to global gene expression and APA changes, and it may explain why the infected individuals have some neuropsychiatric symptoms and long-term brain sequela^[49]^.

The results of APA usage and its correlation with gene expression (Figure 3) could be divided into 4 quadrants. It is reported that about 90% of genes in the long isoform are less stable and produce less abundant proteins, but not in all infected individuals because APA can have different regulatory effects on mRNA stability and translation^[50]^. Genes located in the second quadrant (Figure 3) are related to the short isoform, which is more stable and has higher expression, while those in the fourth quadrant represent the long isoform and are less stable.

As for APP, a variation in the length of the APP 3’UTR, whether long or short, can cause aberrant expression and ultimately lead to a high risk of AD. This may be explained by a study by Mbella EG et al^[38]^. They found alternative polyadenylation of APP mRNA generates two molecules with different 3′UTR lengths and translation efficiencies. The long 3′UTR enhances the translation of APP mRNA compared to the short 3′UTR. Nevertheless, certain proteins from the human brain, CHO cells, and Xenopus oocytes bind only to the short 3′UTR but not the long one^[38]^. These findings suggest a relationship between the effectiveness of mRNA translation and protein binding to the 3′UTR of APP mRNA. The existence of secondary structures in the long nucleotide sequence may be the cause of the absence of these particular interactions between the protein and the long 3′UTR. Either long or short, deviation in length of APP 3’UTR can result in aberrant expression and eventually lead to disease risk.

A total of 267 risk genes for common neurological and psychiatric disorders were found to undergo significant changes in APA following infection, implying that APA could be a potential therapeutic target and predictive biomarker for COVID-19-associated neurological and psychiatric sequela.

Crosstalk between miRNAs and APA are involved in numerous aspects in gene versatility and cell cycle^[51]^. For instance, the gene known as cell division cycle 6 (CDC6) is essential for DNA replication. In mammalian cells, CDC6 has the ability to govern the onset of DNA replication and limit the rate of S-phase entry^[52]^. It has been noted that estrogen can cause the 3’UTR of CDC6 to become shorter. The resulting shortened isoforms can cause aberrant expression of CDC6 by preventing it from being repressed by miRNA. Consequently, research into the potential regulation of miRNAs is very promising. A total of 2404 miRNAs targeting the 408 genes in the four quadrants (Figure 4A, 4B) were predicted using the miRcords, miRTarBase, and TarBase. The miRNAs’ target site information can indicate the impact of APA events and has the potential to be used as a predictive biomarker.

In conclusion, after the SARS-CoV-2 infection, APA usage in neural cells in brain tissue changed in response to viral infection, and the trends are different. Changes in APA are enriched in genes related to RNA splicing, translation, synaptic function, and nervous system function, and these changes are related to neurodegenerative and mental diseases.

## Acknowledgments

The authors would like to thank professor Baofa Sun for his support during the research process.

## Data and Code Availability

The publically available datasets used in this study can be found in GSE159812 (https://www.ncbi.nlm.nih.gov/geo/query/acc.cgi?acc=GSE159812). The data supporting the results in this study are available within the paper and its Supplementary Information. All source datasets are archived at https://www.jianguoyun.com/p/DY-rMxAQtcKtDBjD9r0FIAA. All source codes for the data analysis or figure creation are available at https://github.com/yinggu94/APA.

## Supplemental Figures

Supplementary Figure 1 The cell identity analysis by gene expression.

(A) Each cell type proportion in 7 control individuals. (B) Heatmap plot showing the top 10 most differentially upregulated genes in each cell type identified through unsupervised clustering. Red indicates higher expression; Grey indicates lower expression; the Average expression (avg. exp) scale is shown on the top. (C) Violin plot showing the expression level of cell markers with significantly different percentages per cell type. (D) Heatmap plot showing co-upregulated or co-downregulated gene sets per cluster in RRA. Red indicates up-regulated; Blue indicates down-regulated.

Supplementary Figure 2 Distal PAS Contributions to Neural Function.

Supplementary Figure 3 The Global preference of proximal PAS usage in neurons based on the accumulation rates. Statistical significance is assessed by Kolmogorov-Smirnov test. (A) glutamatergic neurons. (B) GABAergic interneurons. (C) oligodendrocyte. (D) OPC. (E) astrocytes. and (F) microglia.

Supplementary Figure 4 The PAS validation by IGV in each cell type. (A) RPL37A gene in glutamatergic neurons. (B) CIRBP gene in GABAergic interneurons. (C) F3 gene in astrocytes. (D) MT3 gene in oligodendrocytes. (E) B2M gene in OPC. (F) JUND gene in microglial. The same color means that the samples belong to the same group; all the tracks keep the same data range, and the triangle at the bottom indicates the PAS sites.

Supplementary Figure 5 APA usage and its correlation with gene expression level of (A) Astrocyte and (B) Oligodendrocyte.

Supplementary Figure 6 GO terms and pathways of quadrant 1 (upper) and quadrant 3 (lower) genes in different cell types. (A) The network nodes were displayed as pies. Each pie sector is proportional to the number of hits originating from a gene list. The pie charts are colored by cell types, where the size of a slice represents the percentage of genes under the term that originated from the corresponding cell type. Terms with a similarity score > 0.3 are linked by an edge (the thickness of the edge represents the similarity score).

Supplementary Figure 7 Network visualizing the connection between the (A) *CALM1*, (C) *APP*, (E) *ACTB,* and common neurological and psychiatric disorders. The colors of grey, blue, and orange nodes represent targeting miRNAs, genes, and diseases, respectively. The gene expression and APA levels of (B) *CALM1*, (D) *APP*, and (F) *ACTB* in the control group relative to the COVID-19 group.

## Supplemental Tables

Supplementary table 1. PAS metadata.

Supplementary table 2. The top 20 markers of clusters.

Supplementary table 3. PAS changes in neuronal cell types relative to other cell types in control group.

Supplementary table 4. PAS changes in different cell types after infection.

Supplementary table 5. PAS changes in different cell types from SCAPTURE results.

Supplementary table 6. The 408 genes in the four quadrants of all cell types.

Supplementary table 7. The predicted microRNA targeting the 408 genes in the four quadrants of all cell types.

Supplementary table 8. GO terms and pathways of the genes in the four quadrants of all cell types.

Supplementary table 9. The 267 identified risk genes in DisGeNET and BioKA databases and GWAS catalogue.

## Reference

[1] Miura P, Shenker S, Andreu-Agullo C, et al. Widespread and extensive lengthening of 3’ UTRs in the mammalian brain [J]. Genome research, 2013, 23(5): 812–25.

[2] Wang Y, Feng W, Xu S, et al. Extensive Involvement of Alternative Polyadenylation in Single-Nucleus Neurons [J]. Genes, 2020, 11(6):

[3] Gruber A J, Zavolan M. Alternative cleavage and polyadenylation in health and disease [J]. Nature reviews Genetics, 2019, 20(10): 599–614.

[4] Miura P, Sanfilippo P, Shenker S, et al. Alternative polyadenylation in the nervous system: to what lengths will 3’ UTR extensions take us? [J]. BioEssays : news and reviews in molecular, cellular and developmental biology, 2014, 36(8): 766–77.

[5] Parras A, de Diego-Garcia L, Alves M, et al. Polyadenylation of mRNA as a novel regulatory mechanism of gene expression in temporal lobe epilepsy [J]. Brain : a journal of neurology, 2020, 143(7): 2139–53.

[6] Heneka M T, Golenbock D, Latz E, et al. Immediate and long-term consequences of COVID-19 infections for the development of neurological disease [J]. Alzheimer’s research & therapy, 2020, 12(1): 69.

[7] Haidar M A, Shakkour Z, Reslan M A, et al. SARS-CoV-2 involvement in central nervous system tissue damage [J]. Neural regeneration research, 2022, 17(6): 1228–39.

[8] Blake D, Lynch K W. The three as: Alternative splicing, alternative polyadenylation and their impact on apoptosis in immune function [J]. Immunological reviews, 2021, 304(1): 30–50.

[9] Kondrashov A, Meijer H A, Barthet-Barateig A, et al. Inhibition of polyadenylation reduces inflammatory gene induction [J]. RNA (New York, NY), 2012, 18(12): 2236–50.

[10] Zhang X, Chen X, Liu Q, et al. Translation repression via modulation of the cytoplasmic poly(A)-binding protein in the inflammatory response [J]. eLife, 2017, 6(

[11] Jia X, Yuan S, Wang Y, et al. The role of alternative polyadenylation in the antiviral innate immune response [J]. Nature communications, 2017, 8(14605.

[12] Yang A C, Kern F, Losada P M, et al. Dysregulation of brain and choroid plexus cell types in severe COVID-19 [J]. Nature, 2021, 595(7868): 565-71.

[13] Satija R, Farrell J A, Gennert D, et al. Spatial reconstruction of single-cell gene expression data [J]. Nature biotechnology, 2015, 33(5): 495–502.

[14] Li G W, Nan F, Yuan G H, et al. SCAPTURE: a deep learning-embedded pipeline that captures polyadenylation information from 3’ tag-based RNA-seq of single cells [J]. Genome biology, 2021, 22(1): 221.

[15] Zhou Y, Zhou B, Pache L, et al. Metascape provides a biologist-oriented resource for the analysis of systems-level datasets [J]. Nature communications, 2019, 10(1): 1523.

[16] Shannon P, Markiel A, Ozier O, et al. Cytoscape: a software environment for integrated models of biomolecular interaction networks [J]. Genome research, 2003, 13(11): 2498–504.

[17] Ye W, Liu T, Fu H, et al. movAPA: modeling and visualization of dynamics of alternative polyadenylation across biological samples [J]. Bioinformatics (Oxford, England), 2021, 37(16): 2470–2.

[18] Elkon R, Ugalde A P, Agami R. Alternative cleavage and polyadenylation: extent, regulation and function [J]. Nature reviews Genetics, 2013, 14(7): 496–506.

[19] Stroup E K, Ji Z. Deep learning of human polyadenylation sites at nucleotide resolution reveals molecular determinants of site usage and relevance in disease [J]. Nature communications, 2023, 14(1): 7378.

[20] Guvenek A, Tian B. Analysis of alternative cleavage and polyadenylation in mature and differentiating neurons using RNA-seq data [J]. Quantitative biology (Beijing, China), 2018, 6(3): 253–66.

[21] Li J, Liu W, Peng F, et al. The multifaceted biology of lncR-Meg3 in cardio-cerebrovascular diseases [J]. Frontiers in genetics, 2023, 14(1132884.

[22] Balusu S, Horré K, Thrupp N, et al. MEG3 activates necroptosis in human neuron xenografts modeling Alzheimer’s disease [J]. Science (New York, NY), 2023, 381(6663): 1176-82.

[23] Zhang H, Tao J, Zhang S, et al. LncRNA MEG3 Reduces Hippocampal Neuron Apoptosis via the PI3K/AKT/mTOR Pathway in a Rat Model of Temporal Lobe Epilepsy [J]. Neuropsychiatric disease and treatment, 2020, 16(2519-28.

[24] Gaiottino J, Norgren N, Dobson R, et al. Increased neurofilament light chain blood levels in neurodegenerative neurological diseases [J]. PloS one, 2013, 8(9): e75091.

[25] Skillbäck T, Farahmand B, Bartlett J W, et al. CSF neurofilament light differs in neurodegenerative diseases and predicts severity and survival [J]. Neurology, 2014, 83(21): 1945–53.

[26] Kim S, Bai Y, Fan Z, et al. The microRNA target site landscape is a novel molecular feature associating alternative polyadenylation with immune evasion activity in breast cancer [J]. Briefings in bioinformatics, 2021, 22(3):

[27] Afonso-Grunz F, Müller S. Principles of miRNA-mRNA interactions: beyond sequence complementarity [J]. Cellular and molecular life sciences : CMLS, 2015, 72(16): 3127–41.

[28] Xiao F, Zuo Z, Cai G, et al. miRecords: an integrated resource for microRNA-target interactions [J]. Nucleic acids research, 2009, 37(Database issue): D105-10.

[29] Hsu S D, Lin F M, Wu W Y, et al. miRTarBase: a database curates experimentally validated microRNA-target interactions [J]. Nucleic acids research, 2011, 39(Database issue): D163-9.

[30] Sethupathy P, Corda B, Hatzigeorgiou A G. TarBase: A comprehensive database of experimentally supported animal microRNA targets [J]. RNA (New York, NY), 2006, 12(2): 192–7.

[31] Piñero J, Bravo À, Queralt-Rosinach N, et al. DisGeNET: a comprehensive platform integrating information on human disease-associated genes and variants [J]. Nucleic acids research, 2017, 45(D1): D833–9.

[32] Wang Y, Lin Y, Wu S, et al. BioKA: a curated and integrated biomarker knowledgebase for animals [J]. Nucleic acids research, 2023

[33] Hindorff L A, Sethupathy P, Junkins H A, et al. Potential etiologic and functional implications of genome-wide association loci for human diseases and traits [J]. Proceedings of the National Academy of Sciences of the United States of America, 2009, 106(23): 9362–7.

[34] Liu T, Han X, Zheng S, et al. CALM1 promotes progression and dampens chemosensitivity to EGFR inhibitor in esophageal squamous cell carcinoma [J]. Cancer cell international, 2021, 21(1): 121.

[35] Bae B, Gruner H N, Lynch M, et al. Elimination of Calm1 long 3’-UTR mRNA isoform by CRISPR-Cas9 gene editing impairs dorsal root ganglion development and hippocampal neuron activation in mice [J]. RNA (New York, NY), 2020, 26(10): 1414–30.

[36] Hefter D, Ludewig S, Draguhn A, et al. Amyloid, APP, and Electrical Activity of the Brain [J]. The Neuroscientist : a review journal bringing neurobiology, neurology and psychiatry, 2020, 26(3): 231–51.

[37] Tiwari S, Atluri V, Kaushik A, et al. Alzheimer’s disease: pathogenesis, diagnostics, and therapeutics [J]. International journal of nanomedicine, 2019, 14: 5541–54.

[38] Mbella E G, Bertrand S, Huez G, et al. A GG nucleotide sequence of the 3’ untranslated region of amyloid precursor protein mRNA plays a key role in the regulation of translation and the binding of proteins [J]. Molecular and cellular biology, 2000, 20(13): 4572–9.

[39] Ghosh T, Soni K, Scaria V, et al. MicroRNA-mediated up-regulation of an alternatively polyadenylated variant of the mouse cytoplasmic {beta}-actin gene [J]. Nucleic acids research, 2008, 36(19): 6318–32.

[40] Ko J, Mills T, Huang J, et al. Transforming growth factor β1 alters the 3’-UTR of mRNA to promote lung fibrosis [J]. The Journal of biological chemistry, 2019, 294(43): 15781–94.

[41] Douaud G, Lee S, Alfaro-Almagro F, et al. SARS-CoV-2 is associated with changes in brain structure in UK Biobank [J]. Nature, 2022, 604(7907): 697-707.

[42] Huang P, Zhang L Y, Tan Y Y, et al. Links between COVID-19 and Parkinson’s disease/Alzheimer’s disease: reciprocal impacts, medical care strategies and underlying mechanisms [J]. Translational neurodegeneration, 2023, 12(1): 5.

[43] Schou T M, Joca S, Wegener G, et al. Psychiatric and neuropsychiatric sequelae of COVID-19 - A systematic review [J]. Brain, behavior, and immunity, 2021, 97(328-48.

[44] Zhao Y, Shi L, Jiang Z, et al. The phenotype and prediction of long-term physical, mental and cognitive COVID-19 sequelae 20 months after recovery, a community-based cohort study in China [J]. Molecular psychiatry, 2023, 28(4): 1793–801.

[45] Thompson R C, Simons N W, Wilkins L, et al. Molecular states during acute COVID-19 reveal distinct etiologies of long-term sequelae [J]. Nature medicine, 2023, 29(1): 236–46.

[46] Wang R, Lee J H, Kim J, et al. SARS-CoV-2 restructures host chromatin architecture [J]. Nature microbiology, 2023, 8(4): 679–94.

[47] An S, Li Y, Lin Y, et al. Genome-Wide Profiling Reveals Alternative Polyadenylation of Innate Immune-Related mRNA in Patients With COVID-19 [J]. Frontiers in immunology, 2021, 12(756288.

[48] Selinger M, Věchtová P, Tykalová H, et al. Integrative RNA profiling of TBEV-infected neurons and astrocytes reveals potential pathogenic effectors [J]. Computational and structural biotechnology journal, 2022, 20(2759-77.

[49] Boldrini M, Canoll P D, Klein R S. How COVID-19 Affects the Brain [J]. JAMA psychiatry, 2021, 78(6): 682–3.

[50] Wang R, Zheng D, Yehia G, et al. A compendium of conserved cleavage and polyadenylation events in mammalian genes [J]. Genome research, 2018, 28(10): 1427–41.

[51] Sun M, Ding J, Li D, et al. NUDT21 regulates 3’-UTR length and microRNA-mediated gene silencing in hepatocellular carcinoma [J]. Cancer letters, 2017, 410(158-68.

[52] Yan Z, DeGregori J, Shohet R, et al. Cdc6 is regulated by E2F and is essential for DNA replication in mammalian cells [J]. Proceedings of the National Academy of Sciences of the United States of America, 1998, 95(7): 3603–8.

